# Engineering degradation-resistant messenger RNAs in yeast

**DOI:** 10.1101/2020.06.25.169177

**Authors:** Ana L. Franklin, Andrea Macfadden, Jeffrey S. Kieft, Jay R. Hesselberth, Erich G. Chapman

## Abstract

Virally-derived RNA structures provide useful tools for engineering messenger RNA (mRNA) transcripts with unique properties. Here we show that 5’→3’ e<u>x</u>oribonuclease-<u>r</u>esistant RNA structures (xrRNAs) from Flaviviruses can be used to protect heterologously-expressed messenger RNAs (mRNAs) from 5’→3’ degradation in *Saccharomyces cerevisiae* (budding yeast). Installation of xrRNAs ahead of a downstream internal ribosome entry site (IRES) leads to the accumulation of partially-degraded mRNAs designed to undergo cap-independent translation. In a yeast strain favoring cap-independent translation, this strategy led to a 30-fold increase in the enzymatic activity of lysates obtained from cells expressing degradation-resistant reporters. Overall, these finding demonstrate the possibility of coupling multiple viral RNA structures together to enhance protein expression.

## INTRODUCTION

The degree to which the abundance of specific messenger RNAs (mRNAs) in cells correlates with the levels of their encoded proteins is a seminal question in molecular biology.^12^ The abundance of an individual mRNA is determined by both the rate at which it is transcribed and the rate at which it is being degraded. Each mRNA’s unique lifecycle is controlled by many factors including its codon usage^2–4^, translatability^5^, structure of its 5’ and 3’ UTR’s^6–9^ as well as coding regions^10,11^, splice-site selection, choice of polyadenylation signals^12^ as well as its compliment of bound RNA-binding proteins (RBP’s)^13^. In eukaryotes the major pathway of mRNA degradation for RNA pol II transcribed genes takes place through depolyadenylation, dissociation of the 5’-cap binding complex, decapping and ultimately 5’→3’exoribonucleolytic degradation of mRNAs.^14–16^ Recent studies support a model in which mRNA degradation rates are coupled to transcription rates presumably through dynamic association of RBP’s that travel along with transcripts and upon degradation of an mRNA, localize back to the nucleus and associate with promoter regions to effect transcription.^17–20^ Overall, mRNA degradation plays a vital role in regulating gene expression.^21,22^

Viruses take advantage of many aspects of host-cell mRNA biology en route to infecting a cell, becoming translated and undergoing subsequent replication, all the while without being destroyed by the cell’s endogenous mRNA turnover machinery.^23,24^ Several years ago, we described the first three-dimensional structure of a class of 5’→3’ degradation resistant RNA structures in Flaviviruses that we termed xrRNA’s due to their resistance to the 5’→3’ exoribonuclease Xrn1 (**Figure 1**).^25–27^ These conserved RNA structures adopt a “knot-like”, doubly-pseudoknotted structure in which the 5’ end of the xrRNA passes through an encircling 3-helix junction. This creates a topological problem that confounds the exonuclease activity of Xrn1 and other 5’→3’ processive exonucleases.^28^ This property make xrRNA’s useful for designing degradation resistant transcripts in the cell.^29–31^ Highlights of xrRNA’s early applications include the use of xrRNAs in creative 2-color imaging experiments simultaneously depicting transcription, translation and degradation of individual mRNA transcripts^32^, as well as coupling of xrRNA structures to functional small-interfering RNAs (siRNAs)^33^ and “conditional” guide RNAs (cgRNAs)^34^. The experiments described in this manuscript are unique in that the degradation-resistant Xrn1-resistant RNA fragments (xrFrags) produced in this study are able to subsequently undergo translation into functional protein.

**Figure 1:**
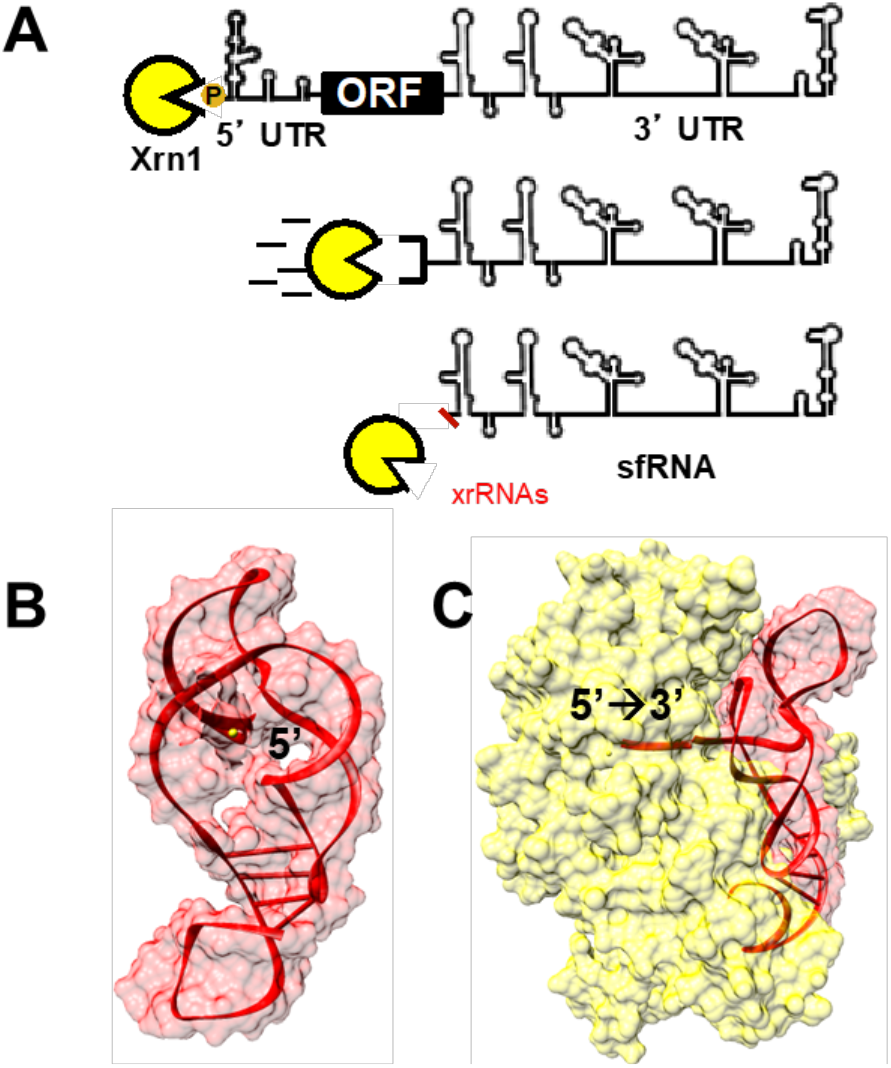
xrRNAs protect downstream sequences from 5’→3’ decay. A) The xrRNA structures examined in this study are derived from the 3’UTR of the Dengue virus where the 5’→3’ blockade of Xrn1 gives rise to <u>s</u>ubgenomic <u>f</u>laviviral RNAs (sfRNAs). B) The three dimensional structures of several xrRNAs have been determined revealing a “knot-like” RNA fold that is unable to be unwound by the helicase activity of Xrn1 and other 5’→3’ exoribonucleases including *E. coli* RNase J1 and yeast Dxo1. C) Docking of Zika virus xrRNA2 and Xrn1. Xrn1 and xrRNAs structures were generated using pdb codes 2Y35 and 5TPY respectively.

Typically, the 5’ m^7^GpppG cap structure that is co-transcriptionally installed on mRNAs transcribed by RNA polymerase II serves as a platform for assembly of the translation initiation machinery. The partially-degraded RNAs that are left following the blockade of Xrn1 lack this structure as Xrn1 is selective for 5’-monophosphorylated RNAs and leaves a 5’ monophosphate upon stalling at xrRNA structures^25^. The design of degradation-resistant mRNAs that remain competent for translation therefore involved using a second viral RNA structure, the internal ribosomal <u>e</u>ntry <u>s</u>ite (IRES) domain from Cricket Paralysis virus (CRPV) which facilitates a streamlined, cap-independent mode of translation initiation that involves structural mimicry of the tRNA-mRNA decoding interaction that takes place in the ribosome.^35–39^ The CrPV IRES has been previously shown to function in a variety of organisms including yeast, highlighting the conservation of the translation machinery across kingdoms.

In this report we demonstrate that coupling of the tandem xrRNA structures found in the 3’UTR of the Dengue virus (DENV) to the CrPV IRES is capable of protecting heterologously expressed mRNAs from 5’→3’ exonucleolytic decay in *Saccharomyces cerevisiae* (budding yeast) and furthermore that the resulting partially-degraded mRNAs serve as effective templates for cap-independent translation originating from the CrPV IRES. The combined effects of the CrPV IRES and xrRNAs produce a significant enhancement in expression of a *lacZ* reporter. Cumulatively, our findings highlight the portability of viral RNA elements and the potential utility of combining functional RNAs from different viruses in to engineer degradation-resistant mRNAs capable of enhanced protein expression.

## RESULTS AND DISCUSSION

### Installation of xrRNA Structures in pre-mRNA Splicing Reporters

To test whether Flaviviral xrRNAs could be used to protect specific mRNAs from 5’→3’ exonucleolytic degradation in yeast we cloned the tandem xrRNA structures from a serotype 2 DENV virus (nt’s 10274-10446 of Genbank accession M20558.1) into a series of previously reported yeast pre-mRNA splicing reporters.^40,41^ As diagrammed in **Figures 2 and 3**, the 2 micron-based reporters are bicistronic. In the parent reporter A (formally pJPS1480), transcription takes place from a constitutively active GPD promoter producing mRNA species with a 91 nt 5’ UTR, followed by exon 1 of yeast ACT1 (ACT1 Orfa), its first intron, and 11 nucleotides of exon 2 (ACT1 Orfb); terminated by a stop codon. The second region of the reporter encodes the intergenic IRES domain from a cricket paralysis virus (CRPV, nt’s 6025 to 6216 of Genbank accession KP974707.1) that is engineered to be in-frame with and drive cap-independent translation of a downstream l*acZ* β-galactosidase reporter. The *lacZ* ORF reporter is trailed by a vestigial copy of the CUP1 gene, then the PGK1 3’ UTR that includes several canonical polyadenylation sites. As diagramed in **Figure 2**, we hypothesized that the inclusion of xrRNA structures directly upstream of the CrPV IRES domain would result in the accumulation of the xrRNA-protected, partially-degraded RNA species. In this model, Xrn1-resistant RNA fragments (xrFrag’s) including the xrRNAs, the IRES domain and *lacZ* ORF persist and remain able to undergo cap-independent translation. As a control we also created a reporter in which simultaneous structure guided mutations to each xrRNA structure render them inoperable (A+2xRNAs_mut_, nucleotides 10303-10305 AGU→UCA and 10377-10379 AGU→UCA). Additional sequence information is included in **Figure 3**.

**Figure 2:**
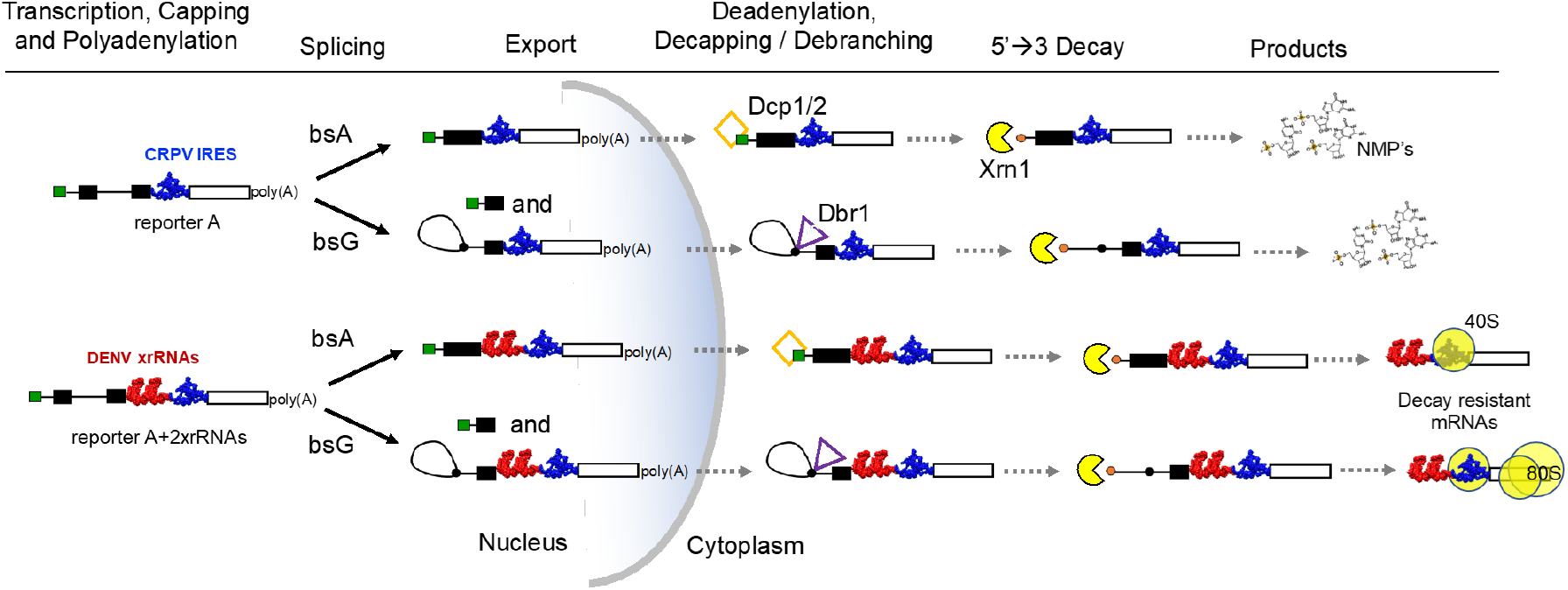
The RNA splicing and decay pathways interrogated by this work. Pre-mRNA splicing reporters were adapted from those created by Mayas, Staley and co-workers in reference ^40^. These reporters are produced from 2 μm plasmids in the nucleus. They undergo pre-mRNA splicing depending on the identity of their branch point nucleotide (bsA or bsG) and are then exported to the cytoplasm where they are eventually subject to decapping and 5’→3’ degradation (top) or <u>d</u>e<u>br</u>anching by Dbr1 followed by degradation by Xrn1.

**Figure 3.**
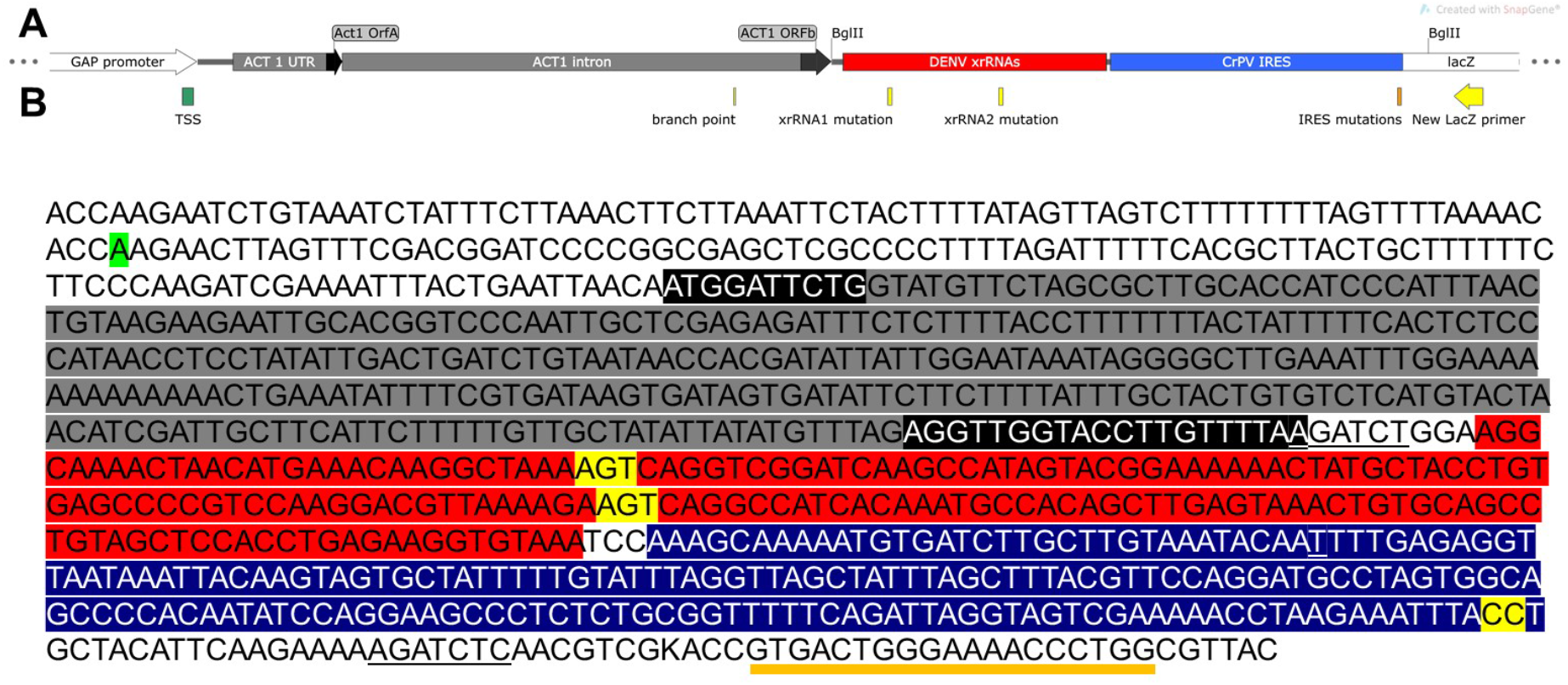
**A)** Layout and **B)** Primary sequence of the intergenic region of the A+2xrRNAs reporter. Features are as labeled within the figure.

### Accumulation of xrRNA-Protected Intermediates

To assess whether partially-degraded RNAs were produced by our reporters A, A+2xRNAs, and A+2xrRNAs_mut_ were transformed into an *imt3 imt4* yeast strain (formerly strain H2545)^40,42^ in which two out of four genomic copies of initiator tRNA_m_ (IMT) genes are deleted. Decreased levels of initiator tRNA_met_ in this system are hypothesized to limit levels of the tRNA_met_-eIF2-GTP ternary complex used in canonical cap-dependent translation.^42^ This yeast strain therefore favors cap-independent translation of the IRES domain included in our experiments.

To assay for the build-up of xrRNA-protected xrFrag intermediates we isolated total RNA and carried out reverse transcription (RT) primer extension (PE) experiments to identify the build-up of 5’→3’ decay intermediates. As shown in **Figure 4**, the inclusion of xrRNA structures results in the build-up of two 5’→3’ truncated, Xrn1-resistant RNA fragments: xrFrag_1_ and xrFrag_2_.

**Figure 4:**
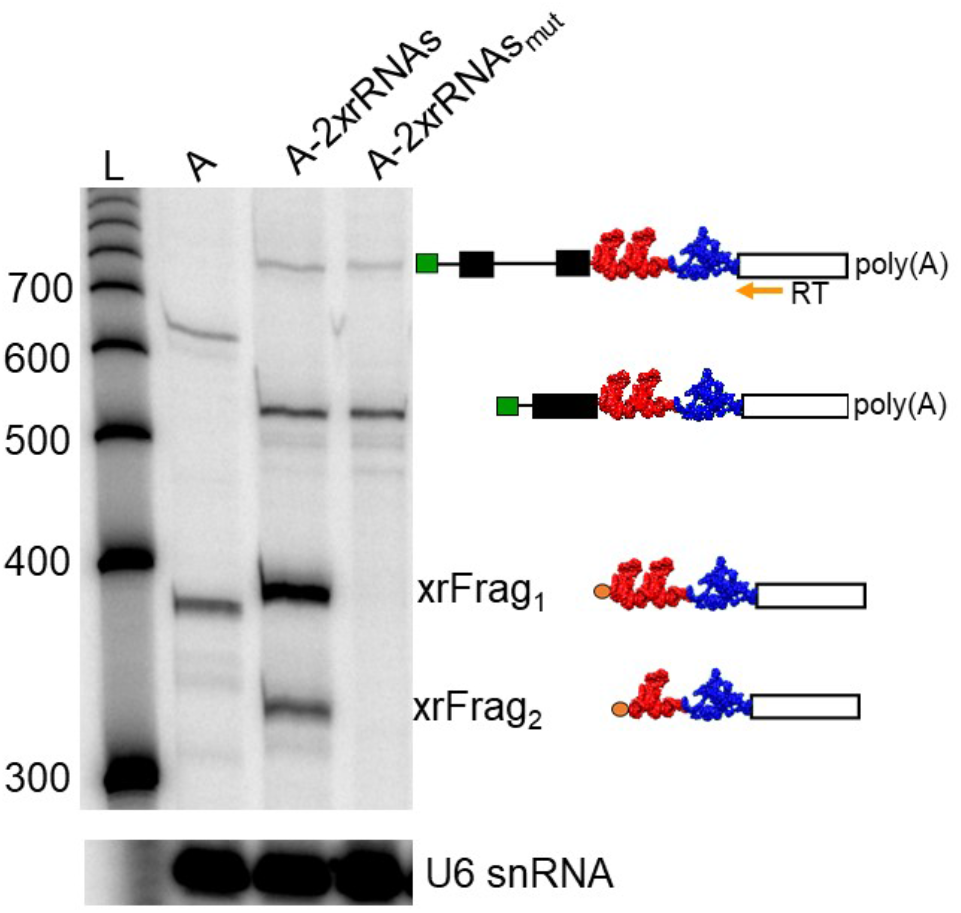
When xrRNA structures are incorporated into mRNAs in yeast partially-degraded RNA species are produced. Primer extension analysis of *IMT3 IMT4* yeast expressing series A reporters. RT was carried out using total cellular RNA and a ^32^P labeled primer complimentary to the 5’ end of the LacZ gene (see figure 3a). The resulting cDNA products were separated by denaturing PAGE. The RNA species portrayed to the right of the figure are those produced from A+2xrRNAs reporter. RT of U6 snRNA served as an internal control.

Specifically, in lane 1 of the gel we observe primer-extension products corresponding to the mRNA expressed by reporter A at 387 nt as well as its un-spliced pre-mRNA precursor at 689nt. In lane 2 we observe similar spliced- and pre-mRNA products for the longer A+2xrRNAs reporter at 560 and 862 nts respectively, as well as two robust bands corresponding to xrRNA-protected fragments xrFrag_1_ and xrFrag_2_ , at 392 and 335 nts respectively. Finally, in the last lane of the gel we observe that a pair of 3 nt point mutations made within in the core of each xrRNA structure disrupt their structure, compromise their decay-resistance and fail to produce partially-degraded RNAs. Cumulatively, these results provide evidence that xrRNA structures can be installed in mRNAs in yeast, fold properly within the cell, don’t interfere with splicing, and function robustly outside of their natural context within the 3’ UTR of the flaviviral genome.

### Translation of xrRNA-Protected RNAs

To detect the predicted increase in protein expression related to the buildup of decay-resistant mRNAs xrFrag1 and xrFrag2, we used a β-galactosidase assay to monitor the enzymatic activity of lysates obtained from the *IMT3 IMT4* yeast cells expressing our series A reporters (**Figure 5a**). When yeast expressing reporter A are lysed by repeated freeze-thaw cycles and analyzed for β-galactosidase activity using ortho-nitrophenyl-β-galactoside (ONPG), we observe OD_420_ values corresponding to 4.3 ± 1.6 Miller units of activity, representing a relatively low level of cap-independent translation occurring from the IRES in transcripts subject to unimpeded decay. When lysates from yeast expressing the xrRNA modified reporter A+2xrRNAs are assessed using the same assay, we measure an average of 127.7 ± 22.0 miller units, an approximately 30-fold increase in enzymatic activity. Yeast expressing reporter A+2xrRNAs_mut_ show β-galactosidase activity similar to the parent construct at 5.0 ± 1.0 miller units.

**Figure 5:**
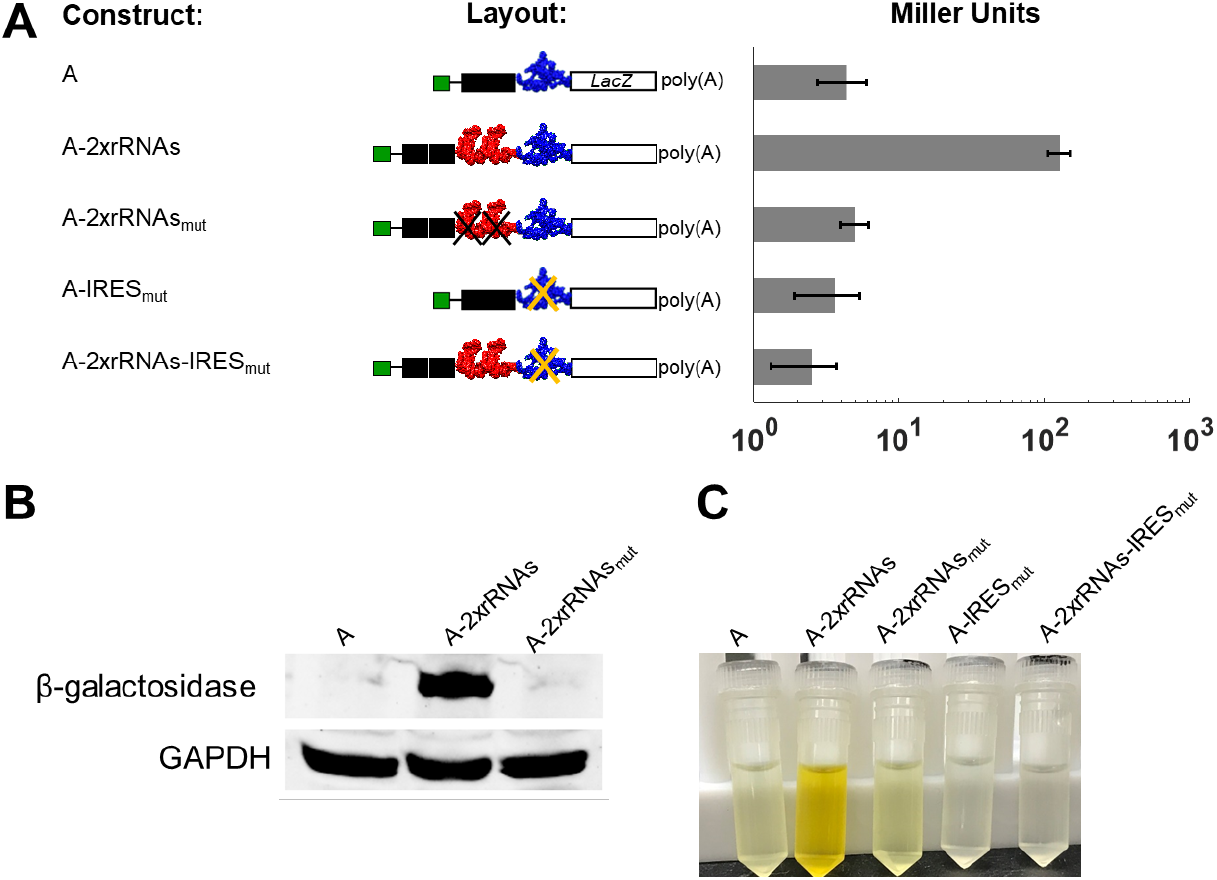
Cap-independent translation of xrRNA-protected reporters increases protein expression relative to unmodified and mutated controls. **A)** yeast expressing A, A+2xrRNAs and A+2xrRNAs_mut_ as well as similar IRES-dead controls A-IRES_mut_, A+2xrRNAs-IRES_mut_, were grown to mid-log phase, lysed and subjected to β-galactosidase assays as described in the Materials and Methods section. **B)** Western blots specific for β-galactosidase and GAPDH from yeast expressing reporters A, A-2xrRNAs, A-2xrRNAs_mut_. **C)** A photo of our samples at the end point of the β-galactosidase assay.

To confirm that the observed increases in β-galactosidase activity were due to increased protein expression, lysates from *IMT3 IMT4* yeast expressing A, A-2xrRNAs, and A-2xrRNAs_mut_ were subjected to quantitative western blotting using a monoclonal β-galactosidase antibody (**Figure 5b**). In our assay we detect substantial levels of β-galactosidase in lysates expressing reporter A+2xrRNAs, though β-galactosidase levels in lysates from yeast expressing A, or A-2xrRNAs_mut_ were below the limit of detection. GAPDH levels remain constant throughout. Combined these data show that *IMT3 IMT4* yeast expressing A-2xrRNAs express substantially more functional β-galactosidase than unmodified (reporter A) or mutated (A-2xrRNAs_mut_) controls.

To establish that translation of the partially degraded reporters is in fact initiated from the CrPV IRES, we also tested two IRES-dead controls. The reporter A-IRES_mut_ (formally pJPS-1482), as well as the analogous xrRNA-modified reporter A-2xrRNAs-IRES_mut_, encode mutations in the IRES domain for two neighboring guanosines that are critical for the formation of a pseudoknot interaction that mimics the tRNA-mRNA codon-anticodon interaction in the ribosome (nt’s G6214→C and G6215→C in the CrPV genome, Genbank accession KP974707.1).^43,44^ Yeast expressing A-IRES_mut_ or A-2xrRNAs-IRES_mut_ produce β-galactosidase activities of 3 ± 1.7 and 2.6 ± 1.2 Miller units respectively suggesting these constructs are not readily translated (**Figure 5a and 5c**). Combined with the previous results that portray the build-up of partially-degraded RNA fragments, these data suggest that the cap-independent translation of the xrFrag’s is responsible for increased protein expression by A-2xrRNAs.

As an means of verifying that translation of partially-degraded, IRES-containing, xrFrag’s takes place, we used sucrose gradient fractionation to isolate polyribosomes from cycloheximide-treated *IMT3 IMT4* yeast. We then used RT-PE to assay for the presence of xrFrag transcripts throughout each gradient. dPAGE analysis of this experiment is included in **Figure 6**. As shown for yeast expressing A+2xrRNAs, we detect primarily xrFrag2 products throughout the gradient together with a shorter, similarly abundant fragment that is harder to assign. The length of this species, ∼250 nt’s, is too short to include either xrRNA, though we speculate that this product could represent Xrn1 stalling at a 40S-IRES complex. Wilusz and co-workers have previously reported stalling of Xrn1 at IRES structures in other viruses.^45^ Furthermore, the *imt3* imt4 yeast strain used in our experiments is specifically engineered to favor the formation of 40S-IRES pre-initiation complexes. Cumulatively, the observation of xrFrag’s associated with polyribosomes and our other results demonstrate that cap-independent translation of xrFrags takes place and leads to significantly more protein production from xrRNA-modified reporters.

**Figure 6:**
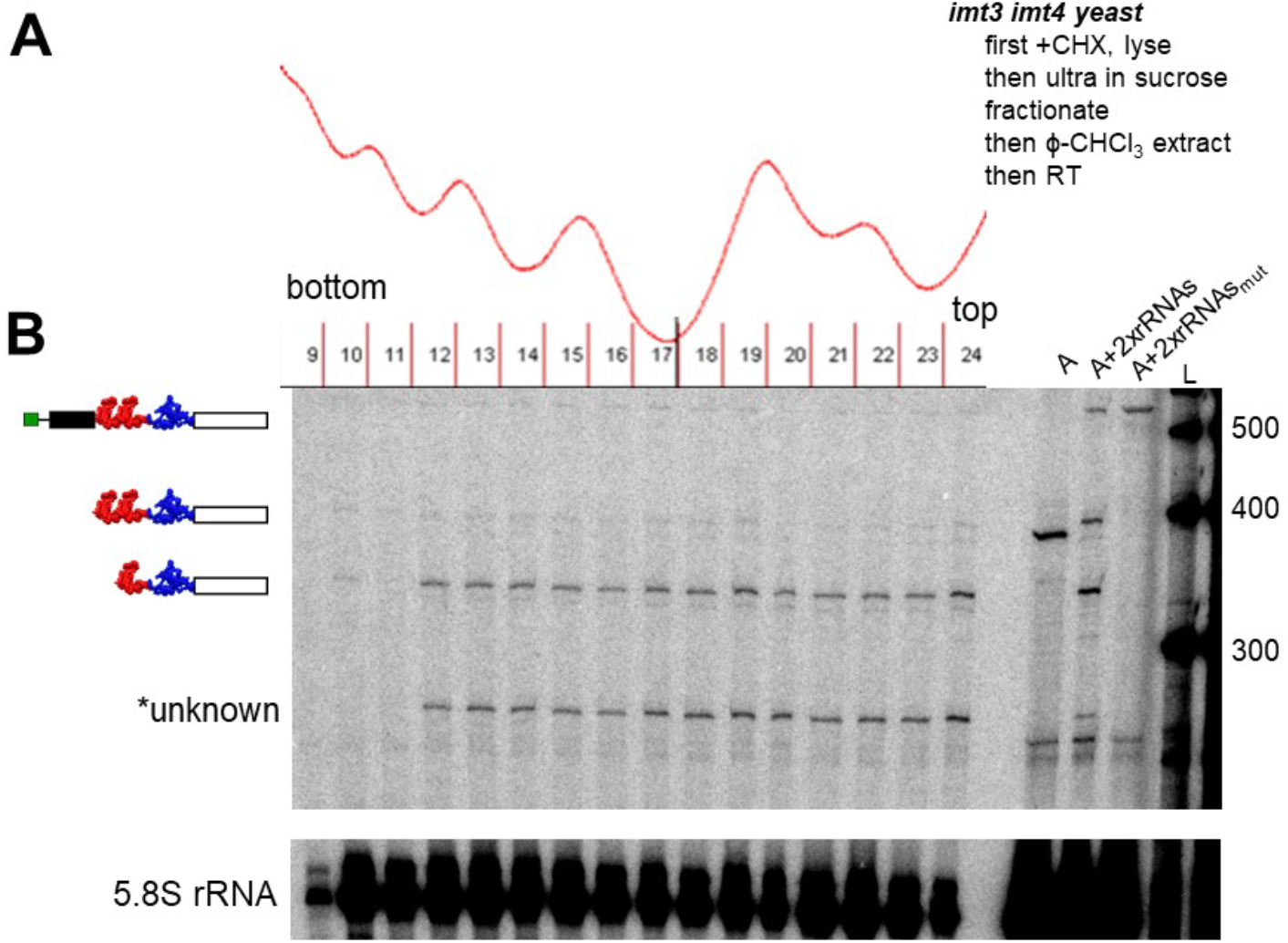
Analysis of polyribosomes from yeast expressing pJPS1481xrRNAs. *IMT3 IMT4* yeast expressing A+2xrRNAs were grown to mid-log phase then translation was arrested using cycloheximide. Yeast were then lysed and fractionated to produce the UV-trace and corresponding fractions shown in part A. Subsequently, individual fractions were phenol-chloroform extracted and then subjected to RT analysis and denaturing PAGE. The resulting gel is displayed in part B. In these experiments RT of the 5.8S rRNA served as an internal control. RT of total cellular RNA of yeast expressing A, A+2xrRNAs and A+2xrRNAs_mut_ is included for reference on the right hand side of the figure as labeled.

### xrRNA Structures Do Not Impede Translation When They Are Installed Within Open Reading Frames

Understanding that xrRNA structures could thus be incorporated in arbitrary transcripts in yeast, we took advantage of the availability of a second set of yeast mRNA reporters^46^ to test whether xrRNAs could be unwound and traversed by the ribosome during translation. Both the ribosome and Xrn1 are obligate 5’→3’ RNA helicases, though operate through distinct biophysical mechanisms.

In order to test whether xrRNA structures can be translated trough, we employed a previously developed dual-luciferase reporter encoding an upstream firefly luciferase (FLUC), a modifiable, in-frame linker region, and a downstream renilla luciferase (RLUC). In these experiments cap-dependent translation of these reporters produces an RLUC-linker-FLUC fusion protein. The influence of specific RNA sequences and tructures, for example by frame-shifting pseudoknots,^46^ on translation is assessed by incorporating them into the linker region and monitoring changes in RLUC versus FLUC activity that result. To assess the influence of xrRNA structures on progression of the ribosome, we generated the seven reporters shown in **Figure 7**.

**Figure 7:**
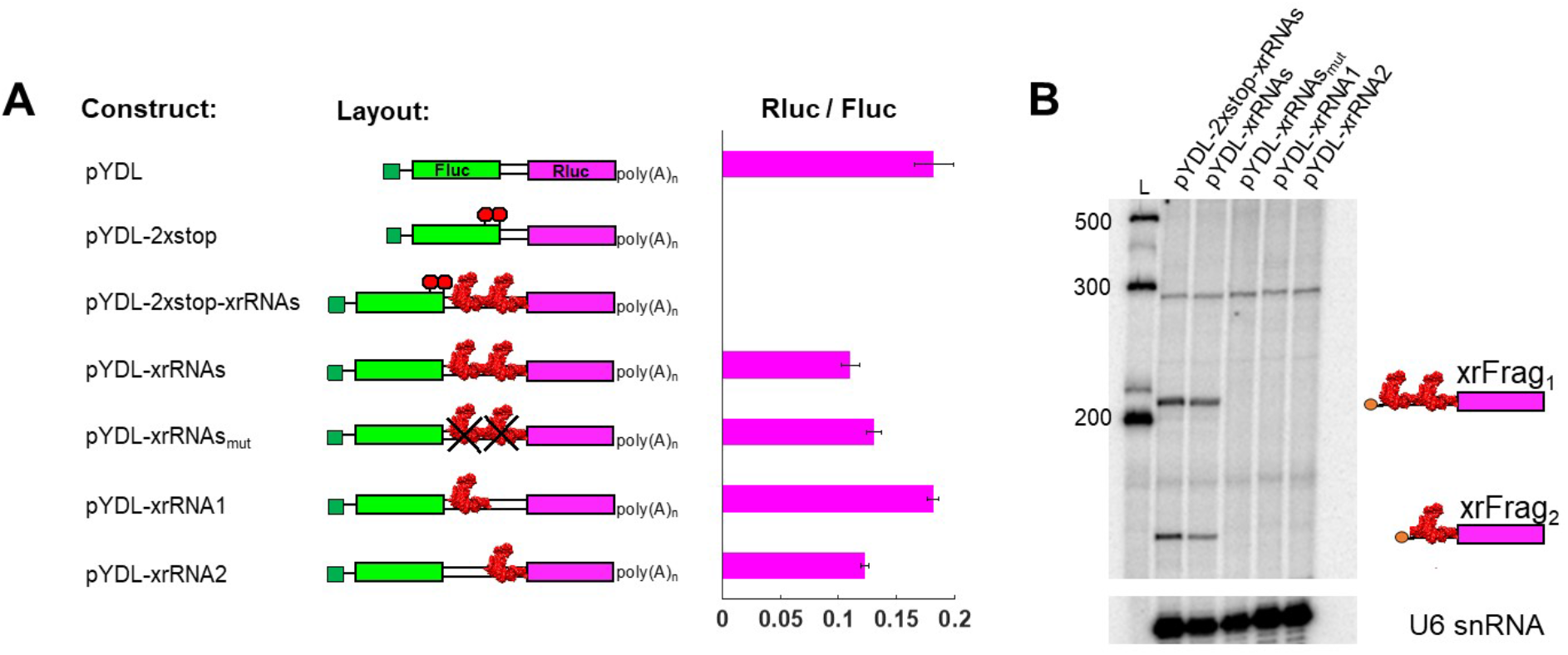
Translation proceeds through xrRNA structures. **A)** The DENV xrRNA structures and related controls were inserted into pYDL dual-luciferase reporters and expressed in yeast. Luciferase assays were preformed using cell lysates to yield the Rluc/Fluc ratios shown next to each construct. B) Total cellular RNA of CRY1 yeast expressing xrRNA-modified pYDL reporters was reverse transcribed and analyzed by denaturing PAGE. Lanes are as labeled within the figure. Interestingly, pYDL reporters that include only a single xrRNA structure fail to produce decay-resistant products. This finding is consistent with other reports that two xrRNA structures are required for production of xrFrag’s *in vivo*.

From top to bottom in **Figure 7** these constructs include: 1) the parent pYDL which produces an RLUC-linker-FLUC fusion protein, 2) a negative control reporter with two stop codons after the FLUC sequence, pYDL-2xStop, 3) an xrRNA-modified version of the previous reporter, pYDL-2xStop-xrRNAs, 4) an xrRNA-modified version of the parent reporter pYDL-xrRNAs, 5) the previous construct with mutated xrRNAs, pYDL-xrRNAs_mut_ 6) the pYDL plasmid with DENVxrRNA1 alone, pYDL-xrRNA1, and 7) and the pYDL plasmid with DENVxrRNA2 alone, pYDL-xrRNA2. The natural DENV xrRNA sequences contain three stop codons that were re-coded by point mutations made in positions listed in the Materials and Methods. These mutations do not compromise the decay resistance of the xrRNA structures themselves (**Figure 7B**).

When yeast expressing these reporters are subjected to dual-luciferase assays we observe a base RLUC/FLUC value of 0.18 ± 0.02 for the parent pYDL reporter. Next, the addition of two stop codons in the linker region proceeding the RLUC open reading frame results in loss of RLUC expression, providing a negative control. Then, as predicted, the incorporation of the tandem DENV xrRNAs behind the stop codon containing control similarly does not produce RLUC activity. For the xrRNA-modified reporter pYDL-xrRNAs, we observe an RLUC/FLUC ratio of 0.11 ± 0.01, a roughly 40% decrease from the parent pYDL construct, though certainly still substantial, indicating that the ribosome can indeed progress through the tandem DENV xrRNA structures. We then explored the translational capacity of several related constructs. The construct with mutated xrRNAs, pYDL-xrRNAs_m_ displayed RLUC/FLUC activity of 0.13 ± 0.01 that is not significantly different than pYDL-xrRNAs providing some indication the three-dimensional structures of the xrRNAs is not responsible for the attenuated RLUC/FLUC values observed when they are incorporated. Installation of DENV xrRNA1 alone in the pYDL linker region produces RLUC/FLUC activity slightly higher than any of the constructs containing DENV xrRNA 2 sequence, including when it is installed alone in pYDL-xrRNA2. This leads us to speculate that perhaps a rare codon within DENV xrRNA2 may underly its attenuated translation, however this hypothesis was not directly tested. Cumulatively, our results demonstrate that the ribosome unwinds and translate through xrRNA elements that conversely, resist being unwound and degraded by Xrn1. These findings portray an interesting difference in how each molecular machine interacts with these unique RNA structures and could be useful in future design of other decay resistant transcripts as well.

## CONCLUSION AND SUMMARY

Experiments in the late 1970’s described the dual role the mRNA cap structure plays in licensing translation and protecting against 5’→3’ decay. Here we used two types of viral RNA elements together to provide these same capabilities in yeast.^47,48^ The observation that RNA structures from divergent positive-strand viruses operate together in an organism that is not a natural host for either of them highlights the conservation of the RNA-decay and translational machinery across domains of life.^49^

This work introduces a new way to stabilize mRNA’s that retain their ability to be translated. Motivation to engineer mRNA’s with enhanced stability and protein coding capacity come exists in multiple sectors ranging from industrial scale enzyme production^50–55^ to current generations of mRNA therapeutics.^6,11,56–61^ Cumulatively, our findings indicate that xrRNA structures present a useful way to prevent the degradation of specific RNA substrates, and when coupled to an IRES element provide a straight forward way to engineer translation-competent xrFrags capable of substantially increasing synthesis of functional protein.

## ACKNOWLEDGEMENTS

The authors thank current members of the Chapman and Kieft labs for critical reading of the manuscript and insightful comments and former members of the Chapman, Hesselberth and Kieft labs for thoughtful discussions and technical assistance. The expression vectors pJPS-1481 and pJPS-1482 were a gift from Dr. Jonathan Staley to JRH. H2545 yeast were a gift from Dr. Sunnie Thompson to JRH. pYDL plasmids were a gift from Dr. Jonathan Dinman to JRH. This work was supported by NIH grants R35GM118070 and R01AI133348 to JSK, R35GM119550 to JRH, and NIH K99GM115757 and R00GM115757 to EGC.

## MATERIALS AND METHODS

### Yeast Husbandry

CRY1 (W303-1A, *MAT*a l*eu2-3*,*112 trp1-1 can1-100 ura3-1 ade2-1 his3-11*,*15*) and H2545 *(MATa trp1-1 ura3–52, IMT1 IMT2 imt3::TRP1 imt4::TRP1 leu2::hisG GAL+)* yeast were maintained using standard yeast husbandry methods. Our recipe for 1 L of synthetic dropout (SD) minimal media includes 2 g synthetic drop-out mix (-His, -Leu, -Trp, -Ura) without nitrogenous base (US Biological cat. D9540), 1.7 g yeast nitrogenous base without amino acids, carbohydrate, or ammonium sulfate (US Biological cat. # Y2030), 5 g of ammonium sulfate, 10 ml of a 4 mg/ml tryptophan solution, 20 ml of an 8 mg/ml leucine solution, 5 ml of a 4 mg/ml histidine, and 865 ml water with 20 g agar added when preparing plates. This solution was autoclaved then supplemented with 100 ml of a sterile filtered 20% (w/v) glucose solution immediately prior to creating liquid cultures or pouring plates.

### Molecular cloning of xrRNA-modified pJPS-1480 series plasmids

Yeast pre-mRNA splicing reporters were modified from reference ^23^. Specifically, the plasmid pJPS-1481 and pJPS-1482 were modified for the studies included here. In order to insert xrRNAs into these constructs, the sequence of the tandem xrRNA structures found in the Jamaica/N.1409 strain, serotype 2 Dengue virus (GenBank accession number M20558.1, nt’s 10272-10448) directly ahead of the CrPV intergenic region (GenBank accession number KP974707.1, nt’s 6025-6216), flanked on both ends by BglII restriction sites were ordered as a gBlock from Integrated DNA Technologies (IDT) and cloned in using standard restriction enzyme-based molecular cloning techniques. Sequences were confirmed by a commercial company using Sanger sequencing.

### Molecular cloning of xrRNA modified pYDL series plasmids

The tandem DENV xrRNAs (specific nt’s listed above) were analyzed for stop codons using an online translation tool (expasy.org). Three stop codons were identified in the natural DENV sequence which were recoded as follows: A10290→T, A10370→T, and T10423→A. Sequences corresponding to the constructs shown in **Figure 4** were purchased as gBlocks from IDT and cloned using Gibson cloning (NEB) into the BamHI site of the pYDL reporter. Specific sequences of any of these constructs is available by request from the authors.

### Transformations into Yeast

Yeast cells intended for transformation, typically 50 ml, were grown to mid-log phase (0.6 ≥ OD_600_≥1.0) pelleted, resuspended and washed once with water, pelleted again and resuspended in 500 µl of 100 mM LiOAc. Cells were transferred to a 1.5 ml tube, again pelleted and finally resuspended in water for use in transformation reactions containing the following: 50 µl resuspended yeast, ∼1 µg of plasmid DNA, 100 µg of sheared salmon sperm DNA, and 500 µl of PLATE buffer (40% PEG 3350, 100 mM LiOAc, 10 mM Tris pH 7.5, and 0.4 mM EDTA). Transformation reactions were incubated at 42 °C for 35 – 40 min then plated onto auxotrophic dropout agar. Plates were incubated at 30 °C for 2 days until a single colony could be picked and maintained for future experimentation and long-term storage.

### RNA isolation

RNA was isolated from yeast by hot acid-phenol extraction. Briefly 10 ml of mid to late logarithmic phase (0.6≥ OD_600_≥ 1.1) culture in Y_min_ -ura was pelleted at 4°C at 3068 rcf. Cells were resuspended in 1 ml of cold Milli-Q grade, autoclaved, filtered water and transferred to screw top 1.5 ml tubes. Cells were pelleted by centrifugation at 6010 rcf in a tabletop microcentrifuge, decanted, and occasionally stored as pellets at -80°C prior to further processing. Pelleted cells were resuspended in 400 µl of TES (10 mM Tris, 1mM EDTA, 0.5% w/v sodium dodecyl sulfate) to which 400 µl of warm 5:1 Acid Phenol:Chloroform (Ambion, cat. # AM9722) were added and the resulting phase-separated mixture was agitated by vortexing at maximum speed for 15 sec. Samples were then incubated in a 65°C heat block for 30 min with periodic vortexing prior to a 5 minute chill on ice and subsequent repartitioning at 18407 rcf at 4°C for 10 min. The aqueous supernatant was transferred to a new 2 ml screw-top tube to which 400 µl of 24:24:1 phenol:chloroform:isoamyl alcohol (Ambion cat. # 9732) was added. The resulting mixture was vortexed for 30 sec and again spun at 18407 rcf for 10 min. The resulting aqueous supernatant was again transferred to a new 2 ml tube and precipitated via the addition of 40 µl of 3 M sodium acetate pH 3.5 and 1000 µl 200 proof ethanol. Samples were incubated on dry ice or in an -80°C freezer for 2+ hours followed by centrifugation at 18407 rcf for at 30 min. The resulting supernatant was decanted and the RNA-containing pellet washed with 70% (v/v) aqueous ethanol prior to re-pelleting at 18407 rcf for 20 min at 4°C in a microcentrifuge. The resulting ethanolic supernatant was decanted and RNA pellets were dried by inversion of the open-capped tubes over a lint-free technical wipe for 30 min to an hour. Upon drying, RNA pellets were resuspended in 50 µl of autoclaved, DEPC-treated, filtered water typically yielding RNA concentrations of ≥2000 ng/µl. Isolated total RNA was diluted and aliquoted to 400 ng/µl, then frozen at -80°C.

### Primer 5’ end-labeling

To label DNA oligonucleotides used for primer extension experiments, oligonucleotides from IDT were received as lyophilized solids were resuspended and diluted to 5 µM. Labeling reactions were designed to use equimolar amounts of DNA and γ-^32^P-ATP (Perkin Elmer cat. # NEG035C001MC) and consisted of 37.5 pmol (7.5 µl of 5 µM) DNA, 40 pmol ATP (1.5 µl of 27.66 µM), and 10 U of T4 PNK in the manufacturer’s supplied conditions of 70 mM Tris-HCl,10 mM MgCl_2_, 5 mM DTT at pH 7.6. Reactions were incubated for 1.5 hrs at 37°C. 150 µl of DEPC-treated water was then added and the reaction passed through a home-poured G-25 sephadex (GE Healthcare cat. # 17-0033-01) spin column. The DNA containing flow through of the procedure was then ethanol precipitated through the addition of 5 µl of high-concentration (>5mg/ml) carrier RNA, 30 µl of 3 M ammonium acetate and 560 µl of 200 proof alcohol. The mixture was incubated on dry ice for at least 2 hrs then pelleted by centrifugation at 21000 rcf at 2°C. The ethanolic supernatant was decanted, and the DNA-containing pelleted washed with 75% ethanol, centrifuged again, decanted, and then dried to completion via speed-vac. Prior to use ^32^P-labeled primers were resuspended in 200 mM Tris pH 8, 200 mM NaCl, 40 mM DTT (6X anneal) buffer.

### Primer extension

Primer extension of total yeast RNA was carried out by combining 12.5 µl of 400 ng/µl total yeast RNA with 2.5 µl of a mixture of ^32^P labeled primers dissolved in 6X anneal buffer. Annealing of ^32^P labeled primers took place in 33 mM Tris, pH 8.0, 33 mM NaCl, 6.67 mM DTT and was accomplished using a thermocycler program that heated to 90°C for 3 min, ramped to 55°C and held for 3 min, then ramped to 4°C. After which the samples were then immediately plunged into liquid N_2_. Prior to reverse transcription samples were thawed on ice then 15 µl of a reverse transcription solution (RT mix) containing 1.6 mM dNTPs, 167 mM Tris pH 8.3, 250 mM KCl,10 mM MgCl_2_, 6 mM DTT, and 1 µl GoScript^TM^ reverse transcriptase (Promega cat. # A5003) was added. The combined mixture was incubated at 42°C for 45 min, 55°C for 15 min, 65°C for 15 min in a thermocycler after which 3 µl of 100 mM NaOH was added and the reaction was incubated for an additional 3 min at 90°C. Lastly, 25 µl of 95% formamide, 5 mM EDTA loading dye was added and the reactions were incubated at 80°C for 3 min prior to being loaded onto a 6% (19:1 mono:bis acrylamide) sequencing gel and electrophoresed at 65 W for 2.5 hrs. The resulting gels were transferred to Whatman paper, dried and exposed to Molecular Dynamics phosphor screens before being imaged on a Storm 720 scanner. Gels were visualized using ImageQuant software.

### *β*-galactosidase assays

Liquid assays were performed as described before. Specifically, 1.5 ml of liquid cultures in SD (-Ura) media were harvested during late logarithmic growth (OD_600_ 0.8–1.2) and washed with Z buffer (60 mM Na_2_HPO_4_, 40 mM NaH_2_PO_4_,10 mM KCl, 1 mM MgSO_4_ pH 7.0). Cells were resuspended in 100 μl of Z buffer and lysed by 6 cycles of freeze-thawing (alternating each 30 seconds between liquid N_2_ and a 42°C water bath). Lysed cells were incubated with 700 μl of prewarmed (30°C) Z buffer that included 1 mg ⁄ ml ortho-nitrophenyl-β−galactopyranoside (ONPG) and 50 mM β-mercaptoethanol (BME). Reactions were incubated at 30°C for 3–4 hrs and stopped by the addition of 0.5 ml 1 M NaCO_3_. Afterwards cell debris was pelleted and the OD_420_ of the supernatant was measured. Activity in Miller units was calculated as OD_420_ × 1,000 / OD_600_ × minutes elapsed × 1.5 ml. Activity in Miller units was corrected for background using lysates from cells lacking the IRES reporter.

### Polyribosome analysis

Yeast expressing series A reporters were grown in -URA synthetic dropout media to an OD_600_ of 0.8, pelleted, and washed twiced with ice-cold YPD supplemented with 0.1 mg/ml cycloheximide. Cells were then washed twice with and resuspended in a lysis buffer consisting of 20 mM Tris-HCl pH 8.0, 140 mM KCl, 1.5 mM MgCl_2_, 0.5 mM DTT, 1% triton X-100, 0.1mg/ml cycloheximide, and 1.0 mg/ml heparin. An aliquot of cold, autoclaved 0.4 mm glass beads was added and cells were alternatingly vortexed for 20 sec and cooled on ice for 100 sec for a total of 5 cycles. Cellular debris and beads were pelleted by centrifugation and the supernatant transferred to a fresh Eppendorf tube, spun down again, and brought to a volume of 1ml using cold lysis buffer. Samples were then layered onto pre-prepared 10-50% sucrose gradients and spun at 35k rpm for 2 hours, 40 min in an SW41 rotor. The resulting gradients were fractionated using a BioComp fractionator with a 0.1mm/sec piston speed, producing ∼5 mm fractions. Individual fractions were then phenol chloroform extracted and ethanol precipitated, quantified and aliquoted as described above.

### Translation Assay

A single colony of yeast transformed with pYDL or pYDL-xrRNA modified reporters was used to inoculate an overnight culture in SD min (-Ura) plus 2% (w/v) glucose media. The following day that overnight culture was used to inoculate a fresh culture which was grown to an OD_600_ of 0.6 to 0.7 units at 30°C. A 1 ml sample of the culture was then removed, pelleted, washed once with phosphate buffered saline (137 mM NaCl, 2.7 mM KCl, 10 mM Na_2_HPO_4_, 1.8 KH_2_PO_4_ pH 7.4) plus protease inhibitors (Roche cat. #04693132001), pelleted again, and resuspended in PBS. Cells were lysed by repeated freeze-thaw cycles as described above for β-galactosidase assays. Cellular debris was then pelleted by centrifugation and either 5 µl or 10 µl of the clarified supernatant used in dual luciferase assays following the manufacturer’s prescribed protocol (Promega, cat # E1910). Data are shown as the ratio of RLUC to FLUC activity averaged over six experiments.

